# Now you see it, now you don’t: optimal parameters for interslice stimulation in concurrent TMS-fMRI

**DOI:** 10.1101/2021.05.28.446111

**Authors:** C. L. Scrivener, J. B. Jackson, M. M. Correia, M. Mada, A Woolgar

**Author notes:** C.L. Scrivener and J.B. Jackson contributed equally to this paper.

## Abstract

The powerful combination of transcranial magnetic stimulation (TMS) concurrent with functional magnetic resonance imaging (fMRI) provides rare insights into the causal relationships between brain activity and behaviour. Despite a recent resurgence in popularity, TMS-fMRI remains technically challenging. Here we examined the feasibility of applying TMS during short gaps between fMRI slices to avoid incurring artefacts in the fMRI data. We quantified signal dropout and changes in temporal signal-to-noise ratio (tSNR) for TMS pulses presented at timepoints from 100ms before to 100ms after slice onset. Up to 3 pulses were delivered per volume using MagVenture’s MR-compatible TMS coil. We used a spherical phantom, two 7-channel TMS-dedicated surface coils, and a multiband (MB) sequence (factor=2) with interslice gaps of 100ms and 40ms, on a Siemens 3T Prisma-fit scanner. For comparison we repeated a subset of parameters with a more standard single-channel TxRx (birdcage) coil, and with a human participant and surface coil set up. We found that, even at 100% stimulator output, pulses applied at least - 40ms/+50ms from the onset of slice readout avoid incurring artifacts. This was the case for all three setups. Thus, an interslice protocol can be achieved with a frequency of up to ~10 Hz, using a standard EPI sequence (slice acquisition time: 62.5ms, interslice gap: 40ms). Faster stimulation frequencies would require shorter slice acquisition times, for example using in-plane acceleration. Interslice TMS-fMRI protocols provide a promising avenue for retaining flexible timing of stimulus delivery without incurring TMS artifacts.

## Introduction

The combination of transcranial magnetic stimulation (TMS) and functional magnetic resonance imaging (fMRI) can be inferentially powerful as it enables direct assessment of how TMS affects neural processing both locally and also in remote, connected brain regions. Moreover, the neural consequences of stimulation can potentially be linked to the impact on behaviour. In the majority of combined studies, TMS occurs before or after the imaging session, outside of the MR environment [“offline”; e.g. 1]. These offline approaches are technically easier to implement than concurrent “online” approaches, where TMS is delivered within the MR scanner. However, offline approaches are not suited for all paradigms, for example, trial-by-trial investigations or where the timing of TMS delivery is critical, and are limited by the duration of the induced TMS effect. Thus, several studies have adopted concurrent methods, for example, to investigate causal influences of higher order brain areas on specialised cortices [2–7]. Recently there has been a resurgence of interest in concurrent TMS-fMRI, for example combining with modern analytical techniques for causal insights into information processing [8, 9]. Current methodological work is improving crucial aspects, such as the signal directly under the TMS coil through development of TMS-dedicated high sensitivity MR coil arrays [10, 11].

One key challenge is that TMS pulse delivery during acquisition adversely affects image quality, producing artifacts [12–14]. These artifacts can be dealt with in several ways. One approach is to remove the affected slices or volumes via temporal interpolation (Fig. 1a) [6, 7, 15]. This can be controlled to result in minimal information loss [16] but is laborious and requires careful assessment to ensure that all affected slices have been removed. Another approach is to use sparse MRI acquisition protocols and interleave TMS pulses with MR readout [e.g. 4, 10, 17]. For example, de Lara et al. [10] delivered TMS pulses in short gaps between MR volumes (Fig. 1b). For their main analysis, they left a gap of 160ms after the TMS pulses, although note that their pilot had indicated a delay of 50ms between the TMS pulse and the subsequence slice was sufficient to avoid artifacts. This approach avoids incurring artifacts and the need to remove associated data. However, interleaving TMS in between volumes reduces temporal resolution and experimental flexibility; for example, stimuli become yoked to the MR acquisition, or gaps have to be very long to allow temporal jitter. Here we examine a third approach, allowing greater flexibility, in which we interleave pulses between fMRI slices (Fig. 1c). This “interslice” approach has been used to implement stimulation frequencies between 1-12.5 Hz [14, 18–23], with the time between the TMS pulse and following slice varying from 34-150ms. However, formal investigation is still required to examine the temporal gap necessary between TMS pulse and slice onset to avoid contaminating the data.

**Figure 1:**
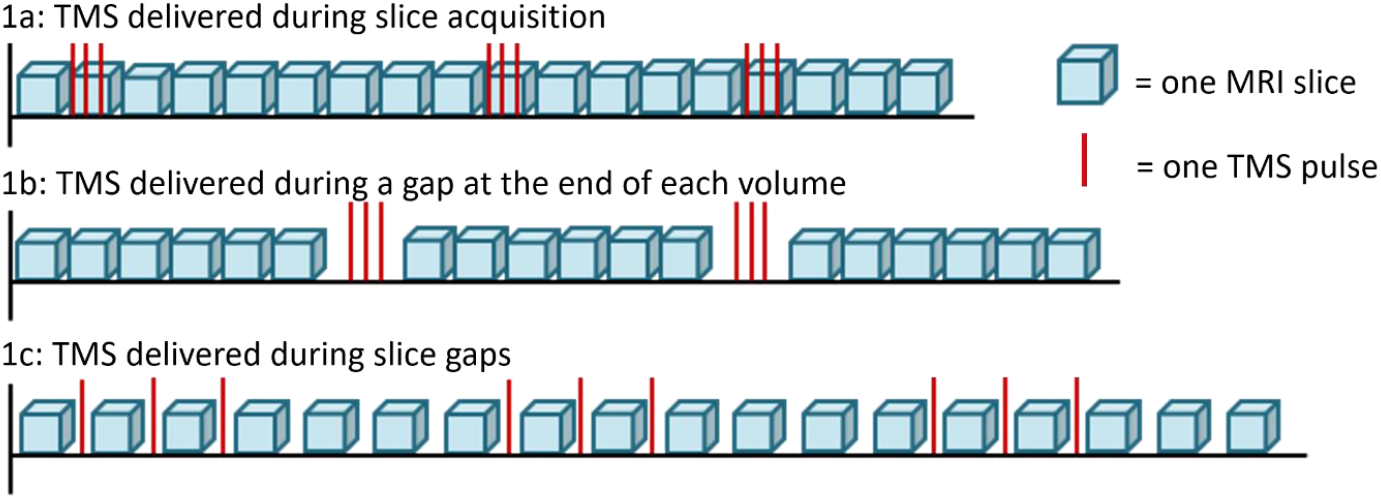
Strategies for concurrent TMS-fMRI. In 1a, TMS is deliberately timed to co-occur with slice acquisition and affected slices are replaced with interpolated data during preprocessing. In 1b, TMS is delivered in temporal gaps between volumes, avoiding any signal loss but reducing experimental flexibility. In 1c, TMS is delivered during gaps between slices, allowing greater flexibility while avoiding the need to remove data.

The aim of this work was to assess the minimum time required between pulse delivery and slice acquisition that could be used without incurring artefacts. We collected data using three setups to compare slice gap duration, MRI coil, pulse timing, number of pulses, and stimulation intensity on data quality.

## Methods

### Data acquisition and Procedure

We acquired FMRI data using a Siemens 3T Prisma-fit MRI scanner in three scanning sessions. In session 1, we used a spherical phantom and two 7-channel TMS-dedicated MR surface coils [11]. In session 2, we recorded from a spherical phantom using a TxRx one channel birdcage coil. In session 3, we acquired with a single participant using the TMS-dedicated surface coils. We selected a representative subset of parameters for sessions 2 and 3 based on the results from session 1.

We delivered TMS pulses using a MagPro XP stimulator and a MagVenture MR-compatible air-cooled figure-of-eight TMS coil. The TMS cable was passed through a wave guide into the MR environment and into the back of the MR bore. During all sessions, the TMS coil was held in position using a coil holder (MagVenture) placed in the bore. In sessions 1 and 3, the TMS coil was mounted on top of one surface coil, and the second surface coil was secured to the coil holder using an in-house custom-built flexible attachment. During session 2, we positioned the TMS coil between the phantom and the TxRx head coil.

For sessions 1 and 3, we acquired functional images using a T2*-weighted echo planar imaging (EPI) acquisition sequence (TE = 30ms, MB factor = 2, no in-plane acceleration, 36 interleaved slices, phase encoding anterior to posterior, transversal orientation, slice thickness 3 mm, voxel size 3 mm x 3 mm, 10% distance factor, flip angle 67 degrees). We increased the minimum possible TR (1.125s) to 2.95s or 1.845s to create temporal gaps of 100ms/40ms between slices.

For session 2, with the TxRx coil, we acquired functional images using a similar T2*-weighted EPI acquisition sequence without multiband acceleration (TE = 30 ms, MB factor = 1, no in-plane acceleration, 18 interleaved slices, phase encoding anterior to posterior, transversal orientation, slice thickness 3 mm, voxel size 3 mm x 3 mm, 10% distance factor, flip angle 67 degrees). We again increased the minimum possible TR (1.125s) to 1.845s to create temporal gaps of 40ms between slices.

We controlled stimulation timing using in-house MATLAB (2014) scripts. A TTL pulse sent from the scanner at the onset of every slice was detected using a National Instruments card. After a set delay determined by the timing parameters, the TMS stimulator was triggered through a BNC connection controlled using a MATLAB toolbox for controlling the Magventure TMS stimulator (created by Michael Woletz, Medical University of Vienna). We tested the fidelity of the timings with this setup with an oscilloscope which recorded sub-millisecond delay and variability in receiving TTL pulses from the MRI scanner and sending subsequent TTL triggers to the TMS machine (see Supplementary Materials for details of timing fidelity).

#### Session 1: Phantom with Surface Coils

We tested the effect of TMS time in relation to slice onset for three amplitudes of TMS (20, 60, 100% MSO) and two EPI sequences (with temporal gaps of 100ms or 40ms). We identified timepoints where TMS resulted in artifacts by sending pulses at timepoints in 10ms intervals from 100ms before (temporal gap; 100ms) or 40ms before (temporal gap; 40ms) to 100ms after the onset of slice readout. We also assessed the cumulative effect of multiple TMS pulses (1, 2, or 3 per volume). For a given set of parameters, all scans were acquired in a single scanner run (e.g. 100% MSO, 1 pulse, across all timings). Each run started and ended with 10 no-TMS (clean) volumes, and with two additional clean volumes in between each timing set (e.g. between TMS at −100ms and TMS at −90ms). We acquired ten volumes for each TMS timing, hitting every volume at the specified time in relation to a predetermined slice. For a single pulse we targeted only slice 9 (slices 3 and 21 in this MB sequence), for two pulses we targeted slices 8 and 9 (including slices 14 and 32), and for three pulses we targeted slices 7, 8, and 9 (including slices 7 and 25).

#### Session 2: Phantom with a One Channel TxRx Coil

We used a TxRx one channel head coil, as this is commonly used for TMS-fMRI, to compare to the data acquired from session 1. We tested one of the parameter sets from session 1 (10ms intervals from −40ms to +100ms in relation to slice onset, 40ms temporal gap, 100% MSO). All scans were acquired in a single scanner run. The run started and ended with 10 clean volumes, and with two additional clean volumes in between each timing set. We acquired ten volumes for each TMS timing, hitting every volume around the acquisition of the 9th slice (slice 18 in this sequence).

#### Session 3: Single Subject with Surface Coils

We performed a final session with a single participant to confirm artifact free data could also be acquired during human TMS-fMRI. The participant gave written informed consent, and data collection was approved by the University of Cambridge Ethics Committee (HBREC.2019.31). We applied TMS 40ms before the onset of slice acquisition, as this time point was artifact free in the phantom data. We applied TMS at 110% resting MT (73% MSO), given that 110% resting MT is commonly used in TMS-fMRI paradigms. Resting MT was determined as the minimum intensity at which a single pulse, positioned over the hand area of the primary motor cortex, produced a visible twitch in the abductor pollicis brevis when at rest, in 10 of 20 successive pulses. We used the MB sequence from session 1 with a 40ms interslice gap, and delivered 1 or 3 pulses per volume (targeting consecutive slices). Each run started and ended with 10 clean volumes, and we acquired 10 TRs with TMS per run. We also included 6 clean volumes between every volume with TMS to give sufficient time between stimulation to comply with TMS safety recommendations [24].

### Analysis

We used two complementary methods to quantify TMS-related image artifacts: tSNR and slice dropout. First, we applied a mask to the EPIs to exclude data from outside the phantom. The mask was created by thresholding data from the auto-align volume to include voxels within the phantom and then binarising it with FSL (v5.0.9). For each parameter set, for each voxel, tSNR was defined as the average signal over its standard deviation through time (FSL).

We quantified dropout as the root mean square deviation (RMSD) of voxels in a slice compared to a mean reference image. For the reference image, we took the average of 10 clean TRs from both the start and end of the scanner run (20 TRs in total). To determine the total possible dropout for each slice, i.e. with no signal recorded, we calculated the RMSD between the averaged clean nifti volume and an empty volume. Average nifti volumes were then created from the 10 TRs in each parameter set (e.g. 10 TRs where TMS was 100%, delivered with a single pulse, at - 100ms in relation to slice onset). To identify the difference in signal caused by TMS, we calculated the RMSD between the clean nifti volume and the TMS volumes. This was then expressed as a percentage of the total possible dropout for each slice, such that 100% dropout indicates that no signal was acquired [following 10]. Calculations were performed using in-house MATLAB (2019a) scripts and the ‘immse’ function.

## Results

### Session 1: Phantom with Surface Coils

Signal dropout results are shown in Figure 2. TMS caused detectable dropout when it was applied at timepoints from 20ms before to 40ms after the onset of slice readout. However, when TMS was applied more than 20ms before, or 50ms or more after the onset of slice readout (i.e., from −100 to −30ms, and from +50 to +100ms) there was no detectable change in RMSD from a clean volume. Dropout was more pronounced with increasing TMS amplitude, but outside of the critical window (−20 to +40ms) there was no detectable effect of TMS, even at 100% MSO. Note that dropout at −20ms was minimal, but to be conservative and due to a slight numerical increase at this timepoint (dropout values in Supplementary Tables 1 to 4), we include it in the critical window. Varying the temporal gap (100ms/40ms) did not change the results.

**Figure 2:**
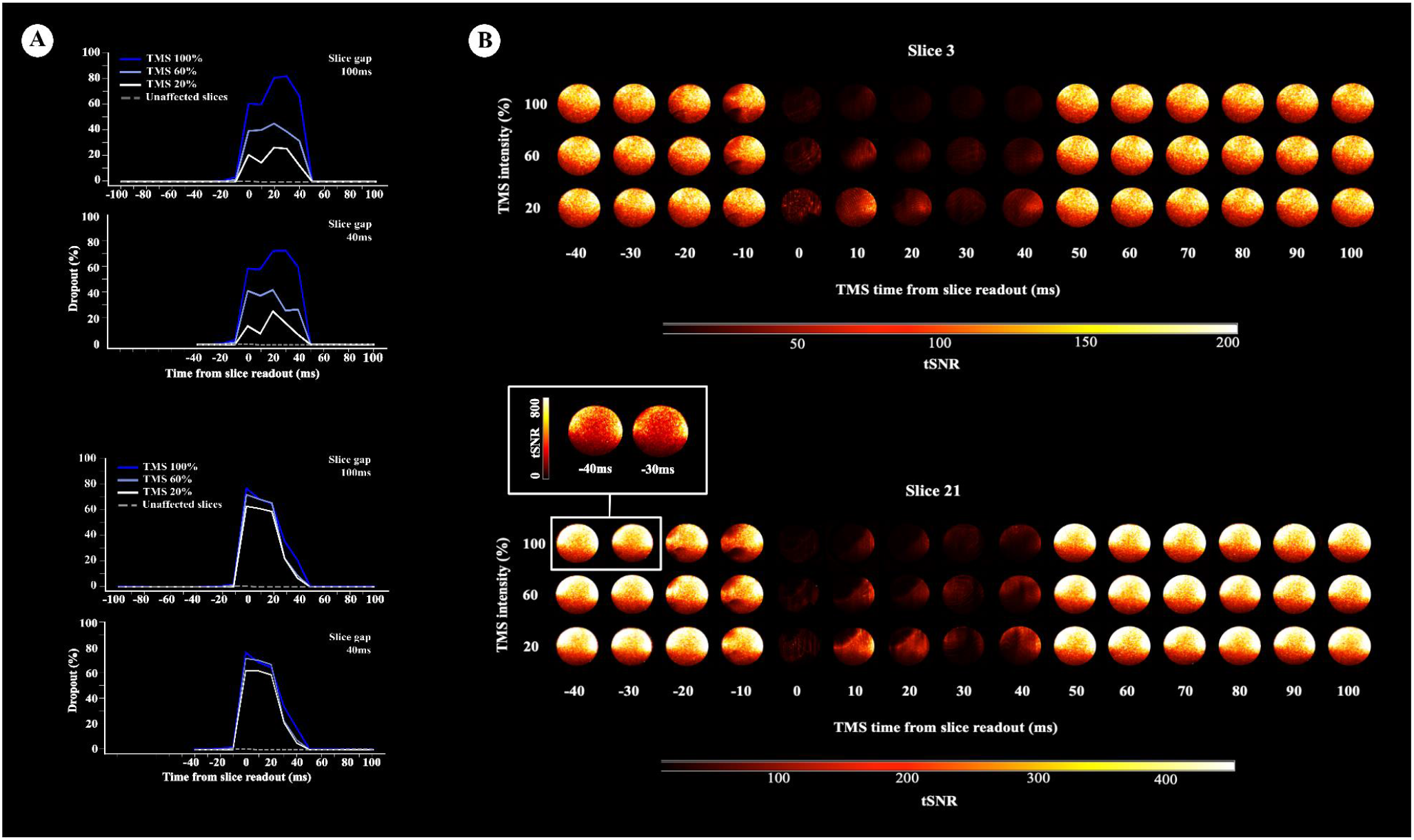
Panel A: dropout in slices 3 and 21 caused by a single TMS pulse applied at different timepoints relative to the onset of slice readout (0ms), at 20%, 60% and 100% MSO. Dropout is expressed as a % of RMSD between the slice with and without TMS, scaled such that 100 % dropout would mean that no signal was acquired (an empty volume). The mean dropout across clean slices (no TMS) is plotted as a comparison in the grey dashed line. Dropout results for both 100ms and 40ms slice gaps are shown. TMS delivered more than 20ms before or 50ms or more after slice onset did not have a detectable effect on dropout. Panel B: modulations in the tSNR caused by a single TMS pulse applied at different time points relative to the onset of slice readout (0ms), using a sequence with a slice gap of 40ms. Two slices per TR were targeted with TMS (slice 3, upper panel, and 21, lower panel), at three TMS amplitudes (20, 60, or 100 % MSO). Note that the maximum tSNR plotted for slice 21 is higher than for slice 3 as it was closer to the MR surface coils. The tSNR results for the sequence with a slice gap of 100ms were very similar, and are not plotted. TMS delivered at 40ms before or 50ms or more after slice onset did not have a detectable effect

We employed a MB acquisition meaning that two slices were affected by each TMS pulse. Slice 3 was near the base of the phantom, and therefore further away from the TMS coil than slice 21. Although the effect of TMS amplitude was more pronounced in the more distant slice 3, the signal in both slices did not differ from baseline (no TMS), when TMS was applied at least 30ms before and 50ms after slice onset.

Visual inspection of the tSNR maps (Figure 2) suggested that small changes in tSNR occurred from 30ms before slice readout, followed by large reductions in tSNR at the onset of slice readout (0ms) that were present until +40ms. However, TMS pulses delivered at least - 40ms/+50ms from slice onset did not modulate tSNR in the affected slices.

If TMS pulses can be delivered at least −40ms/+50ms from slice onset without causing signal loss, then a stimulation frequency of 9.8 Hz is achievable with interslice TMS-fMRI using a slice gap of 40ms and slice time of 62.5ms. We therefore assessed tSNR for sequences where we applied 1, 2, or 3 TMS pulses per volume at 9.8 Hz (Figure 3). At −10ms and at 100% MSO, there was a hint that tSNR reductions might be slightly more pronounced with increasing pulse number, possibly suggesting a cumulative effect of stimulating three slices in a row. However, we do not recommend stimulating at −10ms regardless of this effect, given the increased dropout and reduction in tSNR. When the TMS pulse was applied at −40ms in relation to slice onset, there was no visible effect of the number of pulses on tSNR. It is therefore possible to stimulate multiple times during a single volume in a gap of −40ms before the onset of slice readout, without incurring noticeable artefacts.

**Figure 3:**
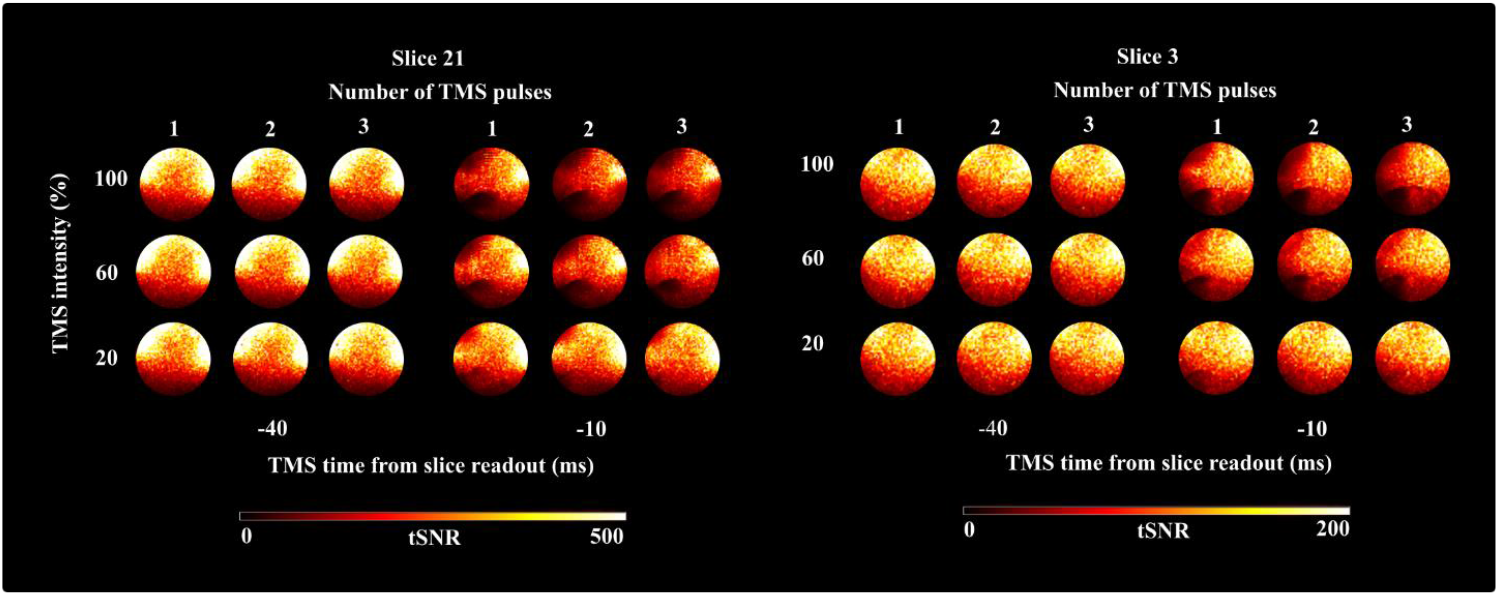
The cumulative effect of multiple TMS pulses (1, 2, or 3 per volume, at 9.8 Hz), at 40ms and 10ms before the onset of slice readout, with a slice gap of 40ms. Two slices per TR were targeted with TMS (slice 21, left, and slice 3, right), at three TMS amplitudes (20, 60, or 100 % MSO). Note that the maximum tSNR plotted for slice 21 is higher than slice 3 as it was closer to the MR surface coils. Small changes in tSNR were found with increasing pulse number at −10ms, particularly in slice 21.

### Session 2: Phantom with a One Channel TxRx Coil

We found the same pattern of results with a conventional TxRx MR coil (Figure 4). Dropout first increased very slightly from baseline at −20ms (see numerical data in Supplementary Table 5) with the largest dropout at 0ms, and returned to baseline at 50ms after the onset of slice acquisition. The tSNR maps also suggested small signal fluctuations from −30ms before slice onset (Figure 4), which was not obvious in the dropout values. In accordance with the results for the surface coils, it was possible to stimulate at −40ms/+50ms from slice onset without incurring signal loss.

**Figure 4:**
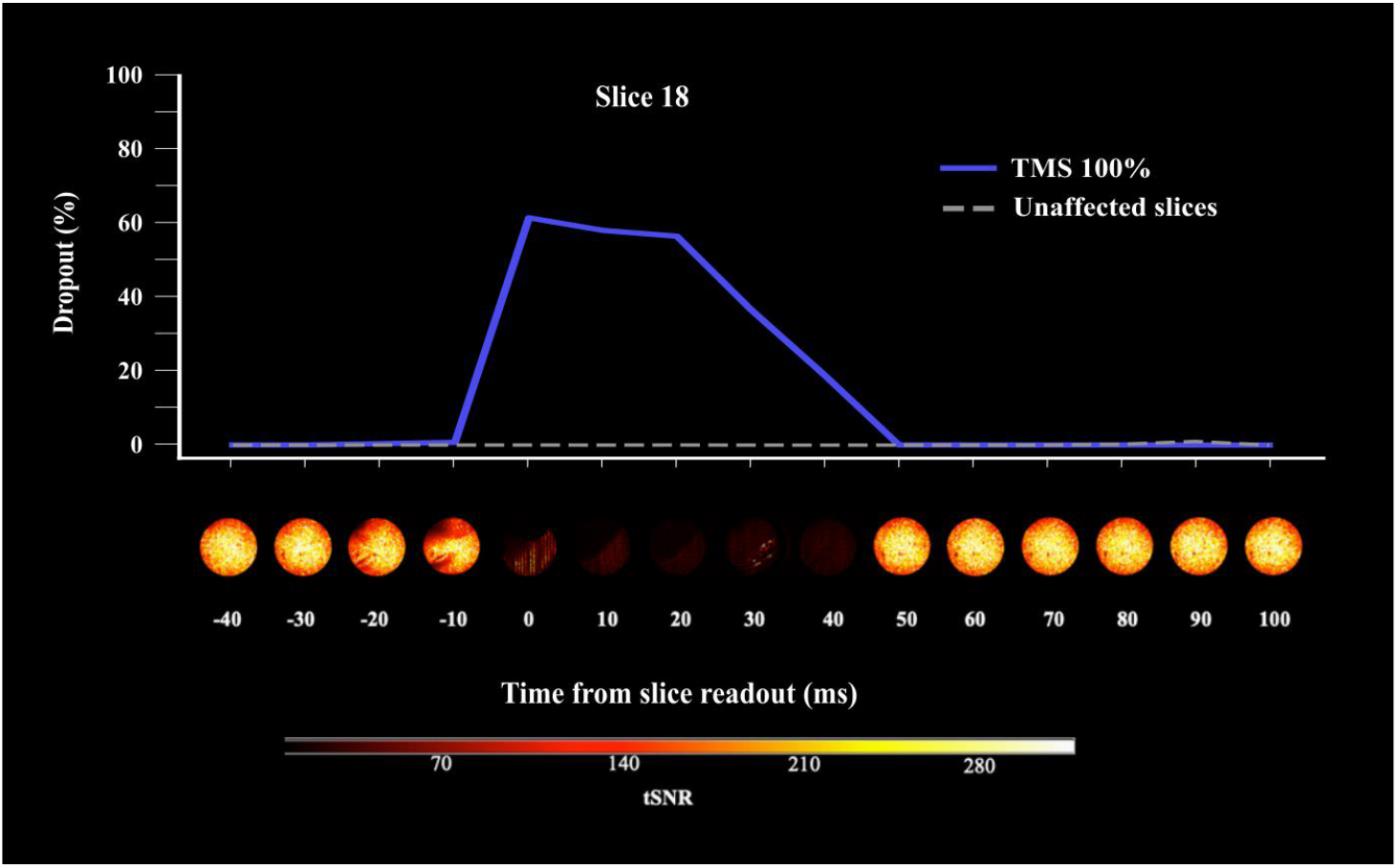
The effect of TMS timing on dropout and tSNR using a TxRx coil. The upper panel depicts dropout caused by a single TMS pulse applied at different time points in relation to the onset of slice readout, where 100% dropout would mean that no signal was acquired. Data was acquired with a TxRx coil and a slice gap of 40ms (and no MB). One slice per TR was targeted with TMS (slice 18), at a single TMS amplitude (100% MSO). The mean dropout across clean slices (no TMS) is plotted as a comparison in the grey dashed line. The lower panel depicts corresponding modulations in tSNR. TMS delivered - 40ms/+50ms from slice onset did not have a detectable effect.

### Session 3: Single Subject with Surface Coils

When a single TMS pulse was applied at −40ms in relation to slice onset, dropout for the targeted slice (21) was 1.31%, which was comparable to the average of 1.23% across all clean slices (Supplementary Table 6). Note that data for slice 3 are not included as this slice was too ventral to be able to assess signal modulations. With three consecutive pulses, dropout for the final targeted slice (21) was 1.12%, compared to an average of 1.70% across all clean slices (Supplementary Table 6). Therefore, the values were very similar across targeted and clean slices, and we do not consider these small fluctuations in signal to reflect artifacts caused by the TMS. As shown in Figure 5, tSNR was also comparable for clean slices to those with TMS applied 40ms before acquisition onset.

**Figure 5:**
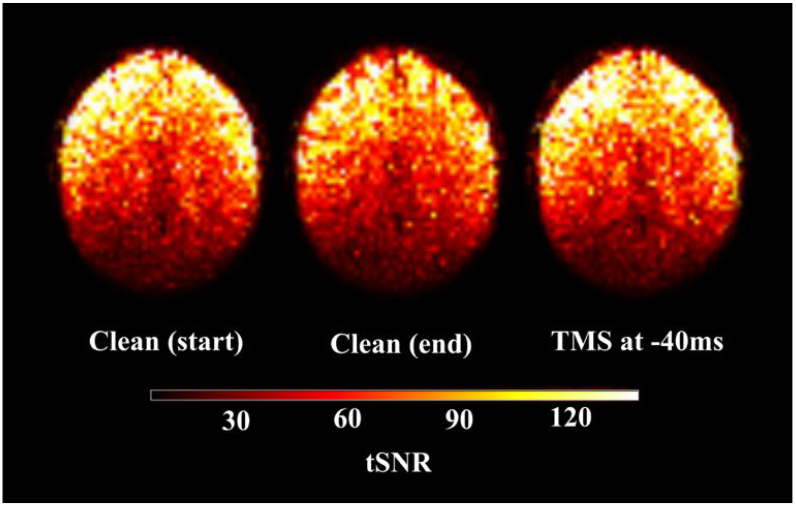
TSNR recorded from a single subject using two surface coils positioned over left and right frontal cortex. Left: tSNR from slice 21 in the absence of TMS (starting volumes). Middle: tSNR in the absence of TMS (end volumes). Right: tSNR in the same slice when a single TMS pulse was applied 40ms before the onset of slice readout, using a sequence with a slice gap of 40ms. The tSNR recorded when TMS was applied at −40ms was comparable to the clean slices with no TMS. The tSNR maps were masked to remove the skull for visualisation.

## Discussion

In the present work, we examined the feasibility of delivering TMS pulses during short gaps between slice acquisitions. We aimed to determine the minimum time required between pulse delivery and slice acquisition to avoid signal contamination. Across all sessions, TMS pulses delivered a minimum of −40ms/+50ms from the onset of slice readout avoided incurring artifacts. This was the case for both MR coils and for pulses delivered at 100% MSO. Thus, an interslice protocol can be achieved for single pulses or trains of TMS with a frequency of up 9.8 Hz using a MB sequence with a slice acquisition time of 62.5ms and interslice gap of 40ms.

As demonstrated here, TMS pulses can be delivered between slices without negatively impacting on data quality. Other approaches time the TMS pulse to coincide with pre-determined slices that are subsequently removed via interpolation. A benefit of this is that no temporal delay needs to be added, either between TRs or slices. Therefore, more data points can be acquired in the same scan duration, increasing the degrees of freedom and statistical power in single-subject analyses [25], as well as the tSNR [26]. However, TMS interference close in time to the radiofrequency excitation pulse can alter the longitudinal magnetisation over several seconds, potentially resulting in false positive activations [14], meaning this timepoint must still be avoided. Moreover, current manufacturers of MR-compatible TMS equipment do not recommend stimulating during slice readout because of theoretical possibility of damaging the MRI machine. As an alternative approach, TMS pulses can be delivered during gaps between TRs, typically lasting up to one second. However, this restricts variation in stimulus-TR onset asynchrony (i.e., the jitter in time between the TMS event and the TR onset) which limits ability to sample the HRF evenly across slices (although this is less of a concern with short TRs). This problem is exacerbated if you need to leave at least 50ms [10] or 100ms [14] between the TMS pulse and TR onset. Instead, delivering pulses between slices facilitates a larger variation in SOA, meaning that the hemodynamic response can be more densely sampled [16, 25].

We showed that interslice TMS-fMRI can also be used in conjunction with repetitive (rTMS), with frequencies of up to 9.8 Hz with the sequence and hardware tested here. If interleaved with slices rather than volumes, the rTMS protocol is not constrained by the duration and timing of the gaps between TRs. Instead, the constraining factor for interslice rTMS is the speed of MR acquisition. Higher rTMS frequencies could be possible with faster sequences but may involve tradeoffs in data quality. While faster sequences enable a higher sampling rate and a relative increase in tSNR [27], they can be associated with additional artifacts [28], and the increase in autocorrelation between samples can lead to false positive effects if not properly accounted for [29, 30].

Interslice protocols have previously been implemented with a range of rTMS frequencies and slice gap durations [e.g. 14, 19-21]. For example, targeting left primary sensorimotor cortex, Bestmann et al. [31] achieved a stimulation frequency of 4 Hz by administering TMS pulses immediately after every slice with a 137ms waiting period prior to subsequent image acquisition (TR 2s, slice duration 113ms). More recently, Hermiller et al. [22] implemented an intermittent theta burst (TR 2.23s, slice gap 107ms), as well as a ‘beta burst’ protocol (12.5 Hz, TR 2.44s, slice gap 34ms), targeting supplementary motor and inferior parietal cortex and reported comparable image quality (using a one-channel head coil). The ability to stimulate at a range of frequencies is necessary for rTMS-fMRI, given that the frequency of stimulation can interact with endogenous oscillations in the targeted region [32, 33]. Thus, if researchers are using a sequence similar to the one tested here (62.5ms slice acquisition time), where an upper limit of ~10 Hz is suggested given possible signal loss, alternative approaches will be necessary for very high frequency paradigms.

In summary, we validated the use of interslice TMS-fMRI with TMS-dedicated surface coils and a standard TxRx birdcage coil. Slice gaps of 40ms were sufficient to prevent degradation of the MRI images, with TMS pulses delivered 40ms before the next slice readout. Therefore, we were able to implement a rTMS interslice protocol of 9.8 Hz without compromising data quality. We recommend the use of this protocol for future studies, but suggest that researchers test their own hardware and sequences. If higher stimulation frequencies are required, then researchers will need to use in-plane acceleration to reduce the time of slice readout, deliver TMS during a delay at the end of the MR volume, or use the temporal interpolation approach, removing the affected slices.

## CrediT Author Statement

Conceptualisation: C.L.S., J.B.J. and A.W.; investigation: C.L.S., J.B.J., M.M.C., M.M., and A.W.; formal analysis: C.L.S. and J.B.J.; writing—original draft: C.L.S. and J.B.J.; writing— review and editing: C.L.S., J.B.J., and A.W.; funding acquisition: A.W.

## Competing Interests

The authors declare no competing interests.

## Data and Code Availability

Raw data and analysis code is available on the Open Science Framework: https://osf.io/tf5wj/

## Acknowledgements

We thank Johan Carlin, Francois Guerit, and MathWorks for their help in optimising the timing fidelity for synchronising MRI and TMS triggers. This work was funded by Medical Research Council (UK) intramural funding SUAG/052/G101400 and Australian Research Council Discovery Project 170101840.

## Supplementary Material

### Timing Fidelity

We confirmed timing fidelity during acquisition by recording the time after each TMS pulse was sent during data collection in session 1 (using the Psychtoolbox function GetSecs). The differences between these timings should be equal to the TR length, given that a TMS pulse was sent at the same time in relation to the TTL received for the same slice for each parameter set. For all recordings with a TR of 2925ms (slice gap 100ms), the mean difference between TMS pulses was 2924.9ms (range 2924.4ms-2925.4ms, S.D. 0.1381ms). For all single pulse recordings with a TR of 1845ms (slice gap 40ms), the mean difference between TMS pulses was exactly 1845ms, (range 1844.3ms-1845.3ms, S.D. 0.0861ms). We also checked the timing of multiple pulses in our repetitive protocol; the time between any two pulses applied to consecutive slices should equal the length of one slice, or 102.5ms. For all recordings with more than one pulse (slice gap 40ms), the mean difference between pulses was exactly 102.5ms (range 101.9ms-102.8ms, S.D. 0.1194ms).

#### Dropout values for session 1: phantom with surface coils

**Table 1:** Dropout in slice 3 caused by a single TMS pulse applied at different timepoints relative to the onset of slice readout (0ms), at 20%, 60% and 100% MSO, using a slice gap of 100ms. Dropout is expressed as a % of RMSD between the slice with and without TMS, scaled such that 100% dropout would mean that no signal was acquired (an empty volume). The mean dropout across clean slices (no TMS) is included as a comparison. The data in the ‘clean slices’ column is the RMSD between slices in the TMS TRs that are not hit with TMS, and the same slices in the clean TRs with no TMS applied. The dropout is always above zero given that the signal fluctuates across TRs. The shaded rows indicate the critical time window during which TMS pulses result in signal dropout. The corresponding plot can be found in Figure 2.

| TMS timing (ms) | TMS amplitude (% MSO) |  |  | Clean slices |
| --- | --- | --- | --- | --- |
|  | 100 | 60 | 20 |  |
| -100 | 0.57 | 0.47 | 0.52 | 0.35 |
| -90 | 0.53 | 0.50 | 0.52 | 0.31 |
| -80 | 0.54 | 0.47 | 0.52 | 0.26 |
| -70 | 0.50 | 0.46 | 0.52 | 0.24 |
| -60 | 0.49 | 0.46 | 0.52 | 0.24 |
| -50 | 0.50 | 0.47 | 0.51 | 0.24 |
| -40 | 0.53 | 0.47 | 0.51 | 0.25 |
| -30 | 0.61 | 0.48 | 0.51 | 0.26 |
| -20 | 1.27 | 0.50 | 0.52 | 0.35 |
| -10 | 4.07 | 1.96 | 0.54 | 0.71 |
| 0 | 60.95 | 39.76 | 21.26 | 0.65 |
| 10 | 60.29 | 40.48 | 15.11 | 0.30 |
| 20 | 80.86 | 45.52 | 26.84 | 0.31 |
| 30 | 82.20 | 39.24 | 26.03 | 0.32 |
| 40 | 66.55 | 31.94 | 12.90 | 0.32 |
| 50 | 0.59 | 0.43 | 0.43 | 0.34 |
| 60 | 0.62 | 0.42 | 0.45 | 0.35 |
| 70 | 0.60 | 0.43 | 0.43 | 0.36 |
| 80 | 0.62 | 0.43 | 0.43 | 0.36 |
| 90 | 0.62 | 0.44 | 0.45 | 0.36 |
| 100 | 0.64 | 0.44 | 0.44 | 0.37 |

**Table 2:** Dropout in slice 21 caused by a single TMS pulse applied at different timepoints relative to the onset of slice readout (0ms), at 20%, 60% and 100% MSO, using a slice gap of 100ms. Dropout is expressed as a % of RMSD between the slice with and without TMS, scaled such that 100% dropout would mean that no signal was acquired (an empty volume). The mean dropout across clean slices (no TMS) is included as a comparison. As a reminder, the data in the ‘clean slices’ column is the RMSD between slices in the TMS TRs that are not hit with TMS, and the same slices in the clean TRs with no TMS applied. The dropout is always above zero given that the signal fluctuates across TRs. The shaded rows indicate the critical time window during which a TMS pulses results in detectable signal dropout. The corresponding plot can be found in Figure 2.

| TMS timing (ms) | TMS amplitude (% MSO) |  |  | Clean slices |
| --- | --- | --- | --- | --- |
|  | 100 | 60 | 20 |  |
| -100 | 0.48 | 0.19 | 0.22 | 0.35 |
| -90 | 0.42 | 0.21 | 0.21 | 0.31 |
| -80 | 0.40 | 0.22 | 0.22 | 0.26 |
| -70 | 0.30 | 0.22 | 0.22 | 0.24 |
| -60 | 0.23 | 0.21 | 0.23 | 0.24 |
| -50 | 0.26 | 0.19 | 0.23 | 0.24 |
| -40 | 0.33 | 0.18 | 0.24 | 0.25 |
| -30 | 0.45 | 0.20 | 0.22 | 0.26 |
| -20 | 0.82 | 0.32 | 0.22 | 0.35 |
| -10 | 2.14 | 1.37 | 0.28 | 0.71 |
| 0 | 77.81 | 72.83 | 63.65 | 0.65 |
| 10 | 69.44 | 69.27 | 61.92 | 0.30 |
| 20 | 65.59 | 66.27 | 59.52 | 0.31 |
| 30 | 36.00 | 22.65 | 22.45 | 0.32 |
| 40 | 21.30 | 9.21 | 7.07 | 0.32 |
| 50 | 0.50 | 0.15 | 0.15 | 0.34 |
| 60 | 0.53 | 0.15 | 0.15 | 0.35 |
| 70 | 0.54 | 0.16 | 0.15 | 0.36 |
| 80 | 0.56 | 0.16 | 0.15 | 0.36 |
| 90 | 0.58 | 0.17 | 0.14 | 0.36 |
| 100 | 0.60 | 0.18 | 0.15 | 0.37 |

**Table 3:** Dropout in slice 3 caused by a single TMS pulse applied at different timepoints relative to the onset of slice readout (0ms), at 20%, 60% and 100% MSO, using a slice gap of 40ms. Dropout is expressed as a % of RMSD between the slice with and without TMS, scaled such that 100% dropout would mean that no signal was acquired (an empty volume). The mean dropout across clean slices (no TMS) is included as a comparison. As a reminder, the data in the ‘clean slices’ column is the RMSD between slices in the TMS TRs that are not hit with TMS, and the same slices in the clean TRs with no TMS applied. The dropout is always above zero given that the signal fluctuates across TRs. The corresponding plot can be found in Figure 2.

| TMS timing (ms) | TMS amplitude (% MSO) |  |  |  |
| --- | --- | --- | --- | --- |
|  | 100 | 60 | 20 | Clean slices |
| -40 | 0.52 | 0.50 | 0.54 | 0.31 |
| -30 | 0.52 | 0.53 | 0.54 | 0.31 |
| -20 | 1.22 | 0.56 | 0.56 | 0.37 |
| -10 | 3.85 | 1.91 | 0.64 | 0.71 |
| 0 | 59.12 | 42.02 | 14.77 | 0.61 |
| 10 | 58.81 | 38.22 | 8.91 | 0.23 |
| 20 | 72.67 | 42.70 | 26.20 | 0.32 |
| 30 | 73.37 | 26.80 | 17.13 | 0.32 |
| 40 | 60.33 | 27.54 | 8.13 | 0.32 |
| 50 | 0.58 | 0.43 | 0.41 | 0.32 |
| 60 | 0.58 | 0.42 | 0.42 | 0.32 |
| 70 | 0.58 | 0.42 | 0.41 | 0.33 |
| 80 | 0.64 | 0.43 | 0.42 | 0.42 |
| 90 | 0.57 | 0.42 | 0.42 | 0.72 |
| 100 | 0.54 | 0.43 | 0.42 | 0.33 |

**Table 4:** Dropout in slice 21 caused by a single TMS pulse applied at different timepoints relative to the onset of slice readout (0ms), at 20%, 60% and 100% MSO, using a slice gap of 40ms. Dropout is expressed as a % of RMSD between the slice with and without TMS, scaled such that 100% dropout would mean that no signal was acquired (an empty volume). The mean dropout across clean slices (no TMS) is included as a comparison. As a reminder, the data in the ‘clean slices’ column is the RMSD between slices in the TMS TRs that are not hit with TMS, and the same slices in the clean TRs with no TMS applied. The dropout is always above zero given that the signal fluctuates across TRs. The corresponding plot can be found in Figure 2.

| TMS timing (ms) | TMS amplitude (% MSO) |  |  | Clean slices |
| --- | --- | --- | --- | --- |
|  | 100 | 60 | 20 |  |
| -40 | 0.42 | 0.21 | 0.23 | 0.31 |
| -30 | 0.37 | 0.23 | 0.22 | 0.31 |
| -20 | 1.07 | 0.35 | 0.22 | 0.37 |
| -10 | 2.03 | 1.18 | 0.30 | 0.71 |
| 0 | 77.20 | 72.48 | 62.90 | 0.61 |
| 10 | 69.46 | 71.01 | 62.61 | 0.23 |
| 20 | 65.40 | 67.71 | 59.47 | 0.32 |
| 30 | 34.18 | 22.96 | 21.47 | 0.32 |
| 40 | 17.29 | 7.79 | 5.34 | 0.32 |
| 50 | 0.51 | 0.14 | 0.16 | 0.32 |
| 60 | 0.52 | 0.14 | 0.15 | 0.32 |
| 70 | 0.53 | 0.14 | 0.14 | 0.33 |
| 80 | 0.59 | 0.17 | 0.14 | 0.42 |
| 90 | 0.55 | 0.17 | 0.15 | 0.72 |
| 100 | 0.53 | 0.16 | 0.16 | 0.33 |

#### Dropout values for session 2: phantom with TxRx coil

**Table 5:** Dropout in slice 18 caused by a single TMS pulse applied at different timepoints relative to the onset of slice readout (0ms), at 100% MSO, using a slice gap of 40ms. Dropout is expressed as a % of RMSD between the slice with and without TMS, scaled such that 100% dropout would mean that no signal was acquired (an empty volume). The mean dropout across clean slices (no TMS) is included as a comparison. As a reminder, the data in the ‘clean slices’ column is the RMSD between slices in the TMS TRs that are not hit with TMS, and the same slices in the clean TRs with no TMS applied. The dropout is always above zero given that the signal fluctuates across TRs. The corresponding plot can be found in Figure 4.

| TMS timing (ms) | 100% MSO | Clean slices |
| --- | --- | --- |
| -40 | 0.39 | 0.33 |
| -30 | 0.38 | 0.34 |
| -20 | 0.74 | 0.35 |
| -10 | 1.09 | 0.37 |
| 0 | 62.00 | 0.36 |
| 10 | 58.59 | 0.35 |
| 20 | 57.01 | 0.37 |
| 30 | 37.11 | 0.34 |
| 40 | 19.23 | 0.33 |
| 50 | 0.39 | 0.35 |
| 60 | 0.38 | 0.34 |
| 70 | 0.37 | 0.35 |
| 80 | 0.38 | 0.55 |
| 90 | 0.36 | 1.34 |
| 100 | 0.35 | 0.32 |

#### Dropout values for session 3: single subject with surface coils

**Table 6:** Dropout caused by a single TMS pulse applied at −40ms in relation to slice readout. Either a single pulse or three pulses of TMS were delivered at 100% MSO, using a slice gap of 40ms.

| TMS parameters | 100% MSO | Clean slices |
| --- | --- | --- |
| -40ms single pulse | 1.31 | 1.23 |
| -40ms three pulses | 1.12 | 1.70 |

## Notes

https://osf.io/tf5wj/

### Competing Interest Statement

The authors have declared no competing interest.

